# Glutathione and thioredoxin systems of the malaria parasite *Plasmodium falciparum*: partners in crime?

**DOI:** 10.1101/128264

**Authors:** Rahul Chaudhari, Shobhona Sharma, Swati Patankar

**Affiliations:** Department of Biological Sciences, Tata Institute of Fundamental Research, Colaba, Mumbai 400005, India; Department of Biosciences & Bioengineering, Indian Institute of Technology Bombay, Powai, Mumbai 400076, India

**Keywords:** *P. falciparum*, Glutathione, Thioredoxin, Antioxidant defense, Antioxidant networks

## Abstract

In *P. falciparum*, antioxidant proteins of the glutathione and thioredoxin systems are compartmentalized. Some subcellular compartments have only a partial complement of these proteins. This lack of key anti-oxidant proteins in certain sub-cellular compartments might be compensated by functional complementation between these systems. By assessing the cross-talk between these systems, we show for the first time, that the glutathione system can reduce thioredoxins that are poor substrates for thioredoxin reductase (Thioredoxin-like protein 1 and Thioredoxin 2) and thioredoxins that lack access to thioredoxin reductase (Thioredoxin 2). Our data suggests that crosstalk between the glutathione and thioredoxin systems does exist; this could compensate for the absence of certain antioxidant proteins from key subcellular compartments.

## Introduction

The malaria parasite *P. falciparum* completes its life cycle in two different organisms: the *Anopheles* mosquito and the vertebrate host. In both hosts, the parasite is subjected to constant oxidative insults. In humans, parasites replicating within red blood cells (RBCs) are bombarded with a high flux of oxygen, resulting in the generation of reactive oxygen species (ROS) [1]. ROS are also generated by metabolic reactions during rapid parasite multiplication, and when the parasite degrades hemoglobin to heme and amino acids [1,2]. Additionally, the apicoplast and the mitochondrion share the heme biosynthetic pathway and electron transport chain, both of which also contribute to ROS production significantly [3-6]. Indeed, parasite-infected RBCs generate twice the amount of ROS compared to uninfected RBCs [1].

Given the susceptibility of parasites to oxidative challenge, efficient ROS detoxification systems are vital. The antioxidant network in the cell is compartmentalized into several key sub-cellular systems within the cytosol, the apicoplast and the mitochondrion [7,8]. These systems achieve detoxification through thiol-disulphide exchange reactions and are comprised of two major branches: the glutathione and the thioredoxin systems. The glutathione system includes glutathione (γ-L-glutamyl-L-cysteinylglycine) as a major regulator of redox homeostasis and other thiol-disulphide oxidoreductases: glutathione reductase (GR), thioredoxin peroxidase-like glutathione peroxidase (TPxGl), glutathione S-transferase (GST), glutaredoxins (Grx) and glutaredoxin-like proteins (GrxlP). The thioredoxin system consists of the small redox-active proteins thioredoxins (Trx), thioredoxin like proteins (Tlp), thioredoxin reductase (TrxR), thioredoxin peroxidase (TPx), peroxiredoxins (Prx) and *Plasmodium*-specific plasmoredoxin (Plrx) [1,2,8].

In accordance with the different subcellular locations of oxidative stress generation, the proteins belonging to each of these systems are also compartmentalized in the parasite. Importantly, some subcellular compartments have only a partial complement of proteins required for thiol-disulphide reactions [7,8]. It is, therefore, likely that functional complementation between the glutathione and thioredoxin systems handles the lack of key anti-oxidant proteins in certain sub-cellular compartments.

To test this hypothesis, we first defined parasite antioxidant pathways and their sub-cellular localizations. This was done by making a comprehensive list of these proteins and their localizations by compiling data from published literature (immunofluorescence and stable transfection of GFP fusion proteins) and data from predictions using online algorithms available on the *Plasmodium* genome database PlasmoDB [9]. The resulting model of the parasite antioxidant network suggested that key sub-cellular locations in the parasite were lacking a few components of the thioredoxin and glutathione systems.

Next, we assessed the possibility of crosstalk between the glutathione and thioredoxin systems by studying the interaction between glutathione and several different thioredoxins using enzymatic assays. For the first time, we show that the glutathione system could play an important role in reducing those thioredoxins that are poor substrates for thioredoxin reductase (Thioredoxin-like protein 1 and Thioredoxin 2) and those thioredoxins that lack access to thioredoxin reductase (Thioredoxin 2). These reactions also could be relevant when thioredoxin reductase is inactivated by electrophilic attack under oxidative stress. Similarly, we show that thioredoxins from the parasite can, in turn, reduce glutathione and act as a backup for the lack of glutathione reductase in a particular subcellular compartment. However, this reaction seems to be of physiological relevance only for Thioredoxin1 and Thioredoxin-like protein 2 as other thioredoxins reduce glutathione disulphide at negligible rates. The results presented in this report suggest that crosstalk between the glutathione and thioredoxin systems exists and that key antioxidant proteins might exhibit flexibility with respect to their reducing partners and their functions.

## Materials and Methods

### Cloning and purification of thioredoxins and thioredoxin reductase from P. falciparum

The *P. falciparum* genome contains genes for three classical thioredoxins (Trx1, Trx2 and Trx3), two thioredoxin-like proteins (Tlp1 and Tlp2) and thiordoxin reductase (TrxR). The cDNA for these were generated using parasite RNA, amplified, and cloned in the bacterial expression vector pET28a (cloning strategy detailed in Supplementary Table 3). All thioredoxins and the thioredoxin reductase were expressed in *E.coli* BL21-DE3 with His-tags. Expressed proteins were purified by Ni^2+^-NTA column chromatography according to the protocol specified by the manufacturer (Ni^2+^-NTA superflow cartridge, Qiagen). Trx2, Trx3 and Tlp2 were always associated with the pellet fraction consistent with the presence of targeting sequences in these proteins. Extensive attempts to improve the solubility of these proteins were not successful. To overcome this problem, we re-cloned these genes lacking the putative organellar targeting signals and found that only Trx2 and Tlp2 could be purified from the soluble fraction (Supplementary Figure 1). The variants of Trx3 were hardly soluble and extensive attempts to improve the solubility of these variants were not successful. After purifying four thioredoxin proteins (Trx1, Trx2, Tlp1 and Tlp2), we systematically tested the activities of these thioredoxins as described in the next section.

### Insulin reduction assay for thioredoxin activity

The activity of each thioredoxin was assayed in an insulin reduction assay that works on the principle of reduction of disulphide bonds in insulin leading to the precipitation of the β-chain [10]. This precipitation can be quantified spectrophotometrically at 600 nm and directly reflects the extent of protein disulphide oxidoreductase activity. Each 500 μl reaction contained 50 mM potassium phosphate buffer (pH 7.4), 1 mM EDTA, 170 μM bovine insulin (in 50mM Tris/HCl, 1 mM EDTA at pH 7.4) and 5 or 10 μΜ of respective thioredoxin. The reaction was initiated at 25°C by adding dithiothreitol (DTT) at 1 mM final concentration. 1 mM DTT with 5 μM bovine serum albumin (BSA) served as a negative control. An increase in absorbance at 600 nm was monitored every 5 seconds for 30 minutes. Activity for each thioredoxin at different concentrations was expressed as a slope of a linear part of a turbidity curve to the lag time (reported as Δ600·min^-2^ × 10^-3^) as described by Martinez-Galisteo *et al* [11]. In additional assays, to identify the functional *Plasmodium falciparum* thioredoxin reductase (PfTR)-thioredoxin pairs, DTT was replaced by 200 μM NADPH and 1 μM PfTR at pH 7.4. In these assays, reaction was initiated by addition of NADPH and increase in absorbance at 600 nm was monitored every 5 seconds for 60 minutes. Activity of each PfTR-Trx pair was expressed as described earlier.

### Thioredoxin reduction by glutathione system

Thioredoxin reduction by the glutathione system was tested using insulin reduction assays. Here, DTT was replaced with glutathione as a reducing agent. Thus, protein disulphide reductase activity of thioredoxins would be observed only if thioredoxins were reduced by glutathione. Each 500 μΙ reaction contained 50 mM potassium phosphate buffer (pH 7.4), 1 mM EDTA, 170 μM bovine insulin (in 50 mM Tris/HCl, 1 mM EDTA at pH 7.4), 10 mM GSH, 1 U·ml^-1^ S. *cerevisiae* glutathione reductase (Sigma) and 10 μM of the respective thioredoxin. The reaction was initiated at 25 °C by adding NADPH at 200 μM final concentration. An increase in absorbance at 600 nm was monitored every 5 seconds for 60 minutes. The reaction without glutathione did not show insulin reduction by any of the thioredoxins, thereby ruling out the possibility of direct reduction of thioredoxins by glutathione reductase. Additionally, assays carried out without thioredoxins confirmed that glutathione alone does not result in insulin reduction. The activity for each thioredoxin at different concentrations was expressed as a slope of a linear part of a turbidity curve to the lag time reported as Δ600·min^-2^ × 10^-3^.

### *In vitro* reduction of GS-SG by purified thioredoxins and thioredoxin reductase

Thioredoxins were tested for their ability to reduce GS-SG. A typical reaction mixture contained 50 mM potassium phosphate buffer (pH 7.4), 1mM EDTA, 5 μM respective thioredoxin, 1 μM PfTR, 250 μM NADPH. Reactions were started at 25 °C by the addition of GS-SG at 1 mM final concentration and the formation of GSH from GS-SG was estimated by following NADPH oxidation at 340 nm. Positive control for the reaction system contained 50 mM potassium phosphate buffer (pH 7.4), 1mM EDTA, 250 μM NADPH, 1mM GS-SG and 0.1 nM GR.

## Results

### A model of anti-oxidant networks in Plasmodium falciparum

The subcellular localizations of antioxidant proteins were predicted by bioinformatics tools and these predictions were compared with experimental evidence as shown in Supplementary Table 1. Most predictions were consistent with the experimental data and enabled us to propose a model network of antioxidant proteins in different compartments of the parasite cell (Figure 1). Although most bioinformatics predictions were consistent with experimental data, a few discrepancies were reported and are listed in Supplementary Table 2a and 2b. After analysis of the discrepancies between predictions and experimental data, we placed nine proteins (Trx2, Trx3, 1-cys-Glrx, AOP, GlP3, TPx_Gl_, GR, TrxR and glyoxalase 2) in all predicted and experimentally defined subcellular compartments. These are shown in a model network of antioxidant proteins (Figure 1) with question marks indicating the discrepancy between predicted localization and experimental localization.

**Figure 1.**
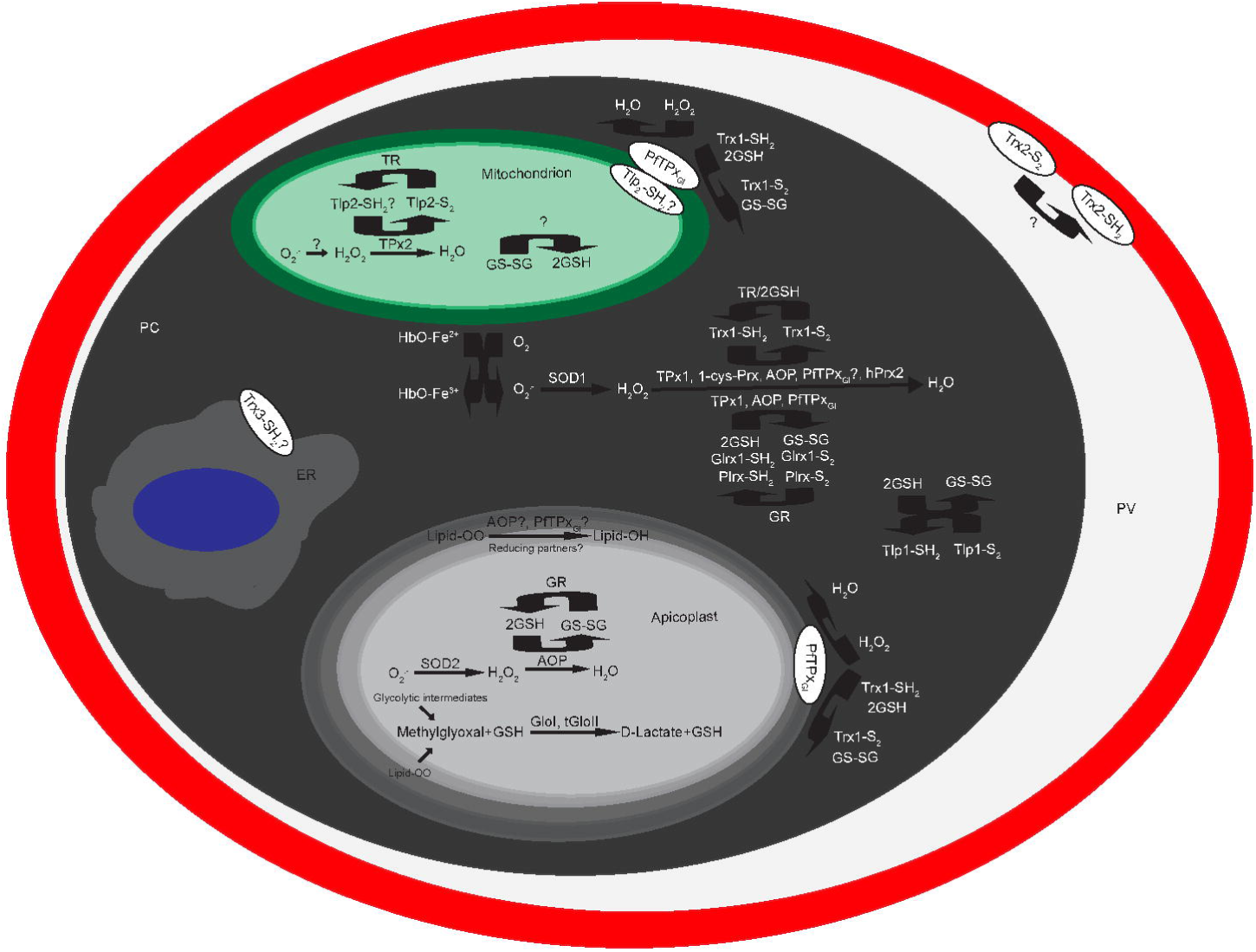
Schematic of the putative model of antioxidant networks in *P. falciparum*. Question marks indicate that the localization is ambiguous or that the reaction proposed is not experimentally verified yet. Abbreviations: HbO-Fe^3+^; oxyhemoglobin with ferriprotoporphyrin, HbO-Fe^3+^; oxyhemoglobin containing ferroprotoporphyrin, O_2_; molecular oxygen, O_2_^-^; superoxide anion, SOD1; superoxide dismutase 1, SOD2; superoxide dismutase 2, H_2_O_2_; hydrogen peroxide, TPx_GI_; glutathione peroxidase-like thioredoxin peroxidase, 1-cys-Prx; 1-cysteine peroxiredoxin, AOP; anti-oxidant protein, TPx1; thioredoxin peroxidase 1, TPx 2; thioredoxin peorxidase 2, Trx1; thioredoxin 1, Trx2; thioredoxin 2, Trx3, thioredoxin 3, Tlp1; thioredoxin like protein 1, Tlp2; thioredoxin like protein 2, Glrx1; glutaredoxin 1, Plrx; plasmoredoxin, TR; thioredoxin reductase, GR; glutathione reductase, GSH; reduced glutathione, GS-SG; oxidized glutathione, Trx-S_2_; oxidized thioredoxin, Trx-SH_2_; reduced thioredoxin, GloI; glyoxalase I, tGloII; targeted glyoxalase II, ER; endoplasmic reticulum, PC; parasite cytosol, PV; parasitophorous vacuole.

This proposed model of antioxidant proteins suggests that a few subcellular locations in the parasite cell lack key components of the antioxidant defense systems. For example, the mitochondrion lacks glutathione reductase. Similarly, the parasitophorous vacuole contains Trx2 as a part of the protein export pump; however, it does not contain thioredoxin reductase required for reducing thioredoxins. Strikingly, the thioredoxin system is completely absent in the apicoplast. The organelle contains neither thioredoxins nor thioredoxin reductase.

The absence of key enzymes like thioredoxin reductase and glutathione reductase in a few subcellular locations suggests that crosstalk between the glutathione and thioredoxin systems might exist to ensure better redox regulation. This led us to ask whether the glutathione and thioredoxin systems can serve as backups for each other.

### Biochemical properties of thioredoxins

Thioredoxins and thioredoxin reductase were purified as described in the materials and methods section. All of these proteins (Trx1, Trx2, Tlp1, Tlp2 and TR), each with a His-tag, were purified to electrophoretic homogeneity as shown by SDS-PAGE (Supplementary Figure 1). Next, the activity of each thioredoxin was assayed in an insulin reduction assay (detailed in the Materials and Methods section). All thioredoxins were found to be active in the dithiothreitol-dependent insulin reduction assays (Figure 2) with varying levels of protein disulphide reductase activity.

**Figure 2.**
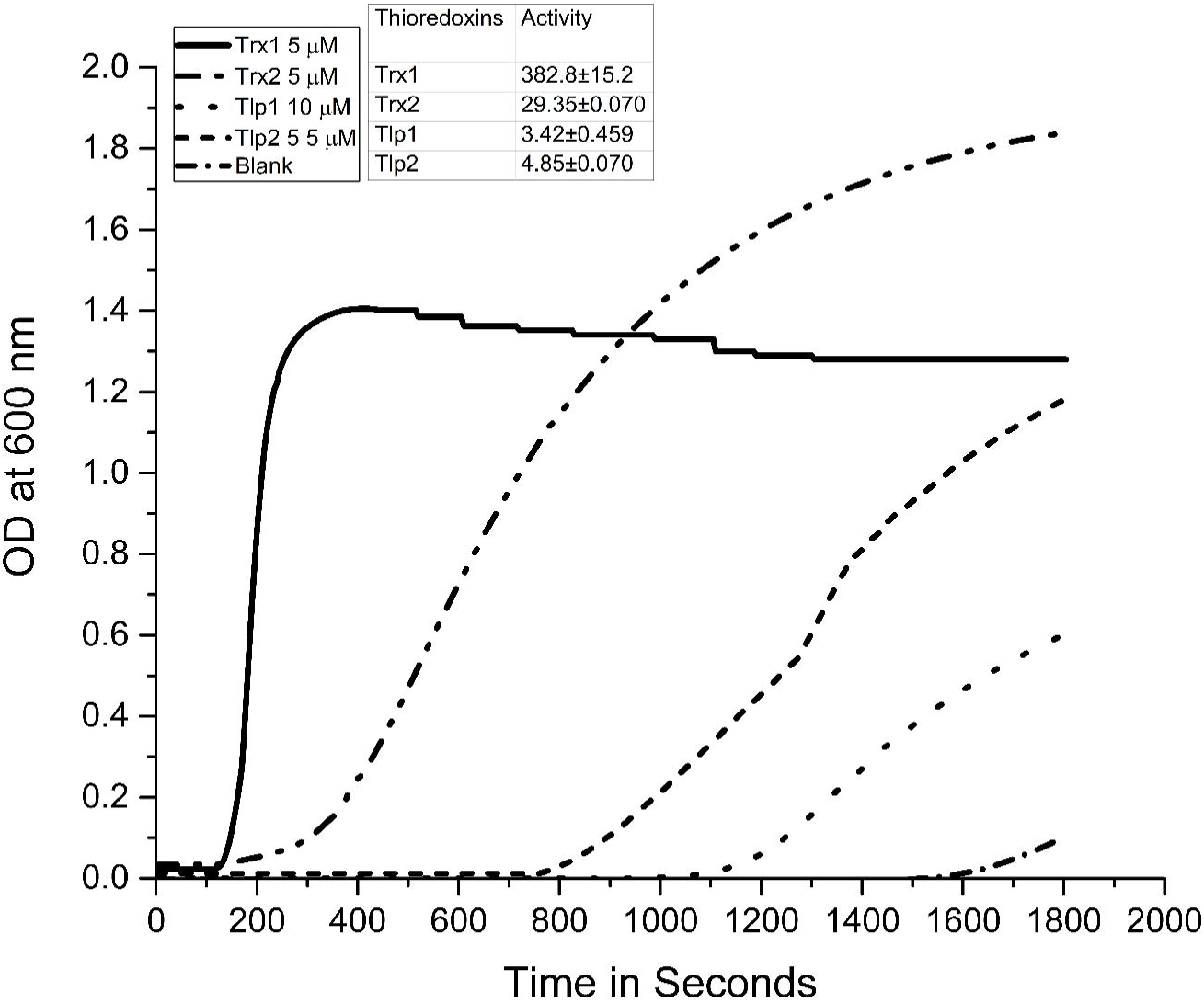
Dithiothreitol dependent activities of different thioredoxins from *P. falciparum* in insulin reduction assay. Turbidity due to thioredoxin mediated insulin precipitation by DTT was measured at 600nm and plotted as function of time. The assay contained 50 mM potassium phosphate buffer (pH 7.4), 1mM EDTA, 170 μM bovine insulin (in 50mM Tris/HCl, ImM EDTA at pH 7.4), 5/10 μM of respective thioredoxin and DTT at 1mM final concentration. 1mM DTT with 5 μM bovine serum albumin (BSA) served as a negative control. Trx1; thioredoxin 1, Trx2; thioredoxin 2, Trx3, thioredoxin 3, Tlp1; thioredoxin like protein 1, Tlp2; thioredoxin like protein 2. The specific activity of each thioredoxin is shown in inset (expressed as Δ600·min^-2^ × 10^-3^ mg^-1^). The data are the mean ± SD of triplicate reactions.

The highest DTT-dependent protein disulphide reductase activity among all thioredoxins was exhibited by Trx1. Trx2 (active site sequence: WCQAC) and Tlp2 (active site sequence: WCAPC) which show partial conservation of the consensus thioredoxin active site motif (WCGPC) showed 12 fold and 75 fold lower activities than Trx1 (active site sequence: WCGPC) (Figure 1 inset). Surprisingly, despite having a consensus thioredoxin active site motif, Tlp1 (active site sequence: WCGPC) at twice the concentration of Trx1, showed 120 fold lower protein disulphide reductase activity than Trx1. The results presented here suggest that thioredoxins might show differential activities under *in vivo* conditions as well.

Next, we examined the ability of *Plasmodium* thioredoxin reductase to serve as an electron donor to these thioredoxins. This was important to analyse as we expected that thioredoxins which are poor substrates of thioredoxin reductase might be linked with other reducing partners *in vivo*. This was tested using insulin reduction assay where TR and NADPH were used as a reducing system. Of the four thioredoxins tested, Trx1, Tlp2 and Trx2 were capable of reducing insulin when incubated with TR. The highest activity was reported for Trx1 followed by Tlp2 (˜4 fold lower activity than Trx1), suggesting that these thioredoxins are linked with TR *in vivo* (Figure 3). Trx2, which demonstrated substantial protein disulphide reductase activity using DTT as a reductant, exhibited far less activity (63 fold and 16 fold lower than Trx1) with TR (Figure 3). Not surprisingly, Trx2 showed less activity with TR. This is expected because Trx2 is located in the parasitophorous vacuole where there is no TR. Additionally, Tlp1 did not show any activity with TR as a reducing partner. These results suggested that Trx2 and Tlp1 might be associated with other reducing partners *in vivo*.

**Figure 3.**
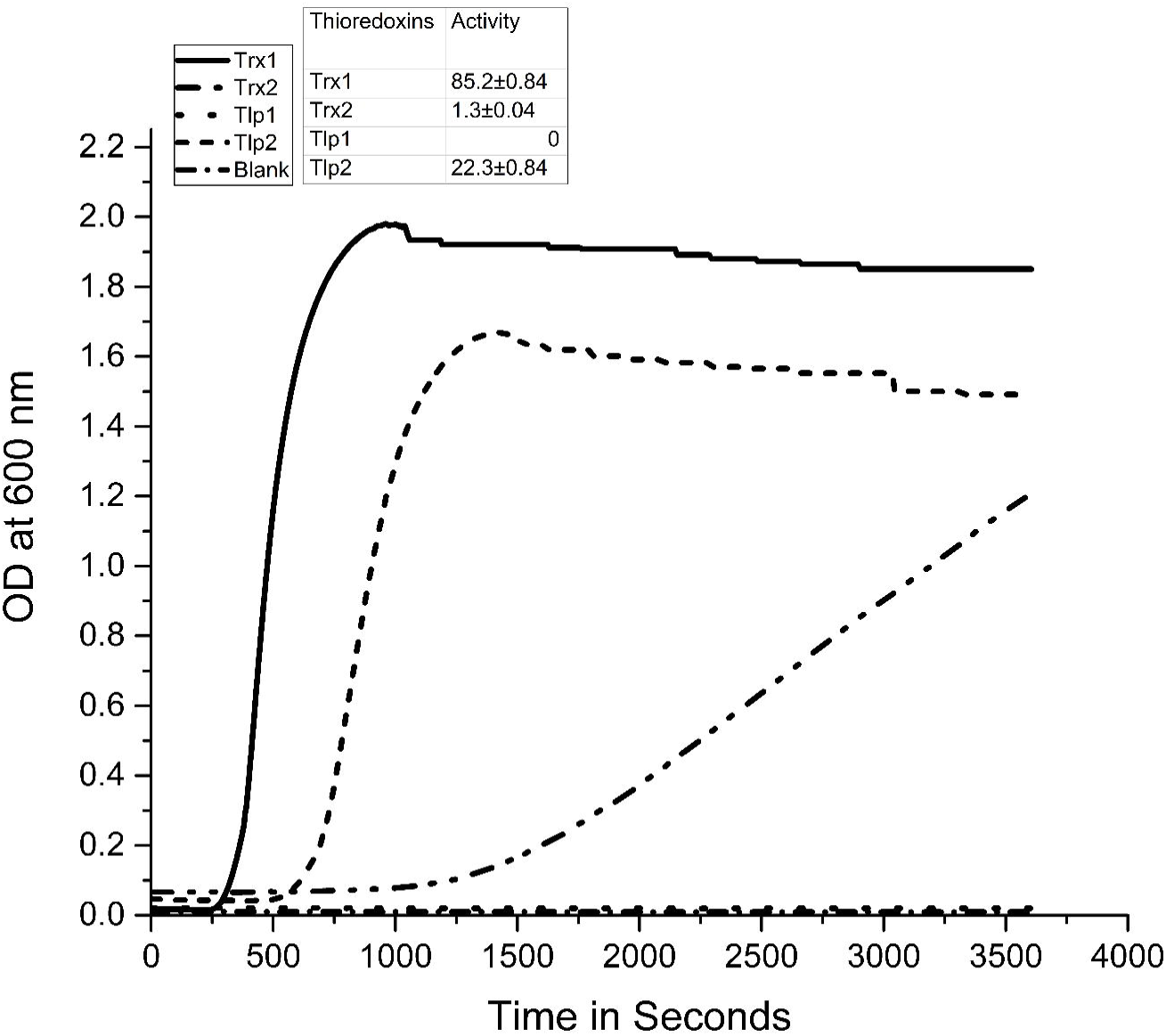
Comparison of Insulin reduction by *P. falciparum* thioredoxin system components. Turbidity due to insulin precipitation by thioredoxin system components was measured at 600nm and plotted as function of time. The assay contained 50 mM potassium phosphate buffer (pH 7.4), 1mM EDTA, 170 μM bovine insulin (in 50mM Tris/HCl, ImM EDTA at pH 7.4), 10 μM of respective thioredoxin, 200 μM NADPH and 1 μMPfTR. The reaction without any of the thioredoxins served as a negative control. Trx1; thioredoxin 1, Trx2; thioredoxin 2, Trx3, thioredoxin 3, Tlp1; thioredoxin like protein 1, Tlp2; thioredoxin like protein 2. The specific activity of each thioredoxin is shown in inset (expressed as Δ600·min^-2^ x 10^-3^ mg^-1^). The data are the mean ± SD of triplicate reactions.

### Only Trx1 and Tlp2 support GS-SG reduction in the parasite

In a possible interaction between glutathione and thioredoxin systems, *P. falciparum* Trx1 has been shown to reduce GS-SG *in vitro*. This reaction seems to be relevant for a particular subcellular location that lacks glutathione reductase or under conditions of enzyme insufficiency. Therefore, we decided to test whether any of the other thioredoxins could reduce GS-SG *in vitro*. This was investigated by assessing the reduction of GS-SG in presence of purified TR, NADPH and each of the four different thioredoxins. In this assay, *E. coli* glutathione reductase was used as a standard for GS-SG reduction.

The data from Figure 4 indicates that TR cannot reduce GS-SG in the absence of Trx1 or Tlp2. The Trx1-TR and Tlp2-TR combinations reduced GS-SG with different efficacies. Other thioredoxins, viz. Trx2 and Tlp1, showed GS-SG reduction at negligible rates. These results suggest that, in the malaria parasite, GS-SG reduction by thioredoxins might be physiologically feasible only in the cases of Trx1 and Tlp2.

**Figure 4.**
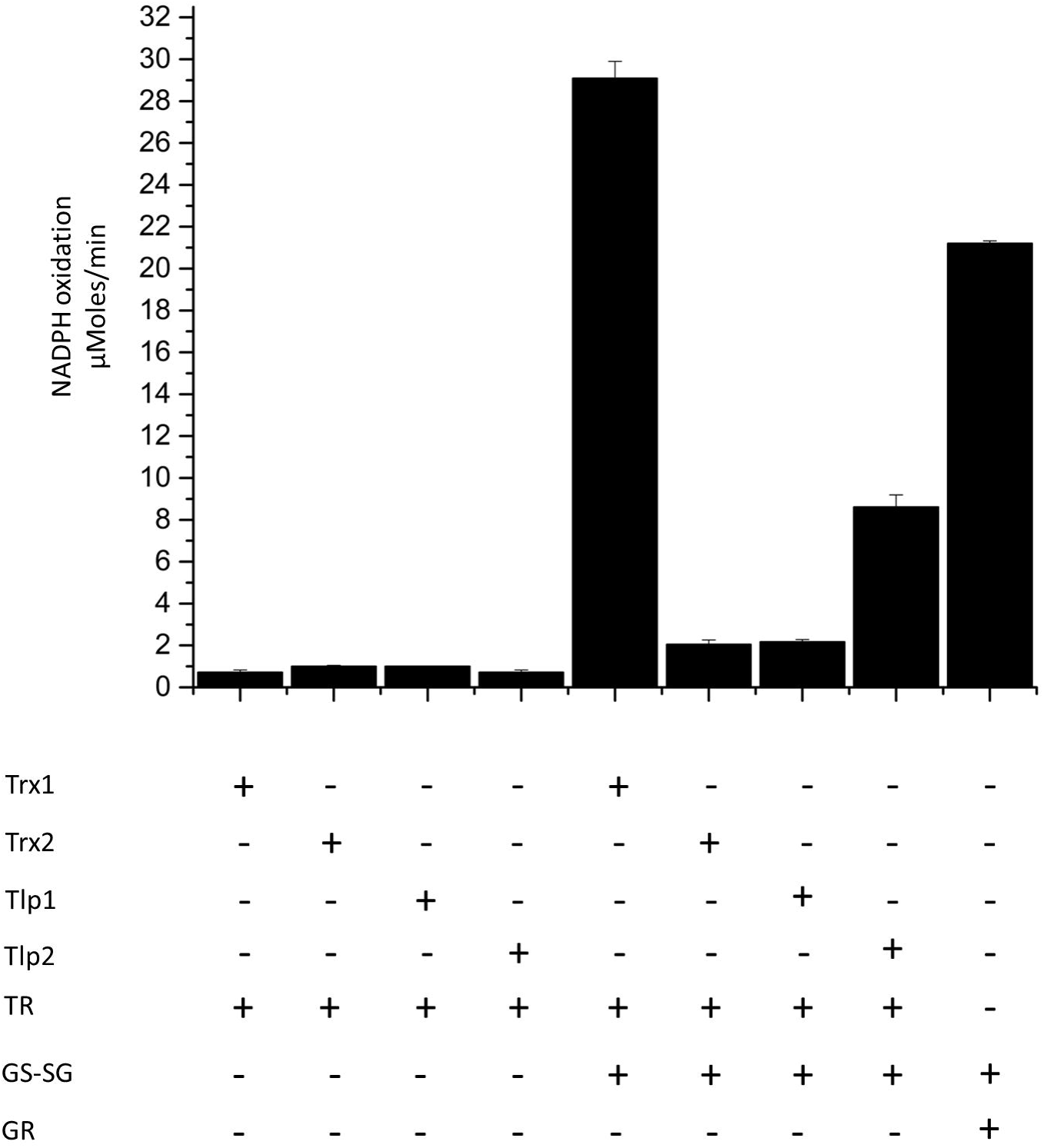
*In vitro* GS-SG reduction by *Plasmodium* thioredoxin system components. Only Trx1 and Tlp2 support reduction of GS-SG *in vitro*. Purified thioredoxins were reconstituted with the indicated components and the NADPH consumption was monitored at 340 nm. Error bars indicate S.D. of three replicates.

### Glutathione can reduce Plasmodium thioredoxins which are poor substrates of TR

*Plasmodium* thioredoxins are localized to different subcellular compartments (Figure 1). Some of these compartments lack thioredoxin reductase, thereby questioning the presence of a complete functional thioredoxin system at those locations. Additionally, when we checked for functional TR-thioredoxin pairs, we found that Trx2 was a poor substrate for TR and that Tlp1 could not serve as a substrate at all.

In order to understand alternative mechanisms for the reduction of these thioredoxins, we performed biochemical assays using Trx-dependent insulin reduction. In these assays, DTT was replaced with glutathione as a reducing agent. Therefore, protein disulphide reductase activity of thioredoxins would only be observed if the thioredoxins were reduced by glutathione. This was verified by performing the assay without glutathione, in whose absence insulin reduction by any of the thioredoxins did not take place. This ruled out the possibility of the direct reduction of these thioredoxins by glutathione reductase, and not by glutathione itself. Additionally, assays carried out without thioredoxins confirmed that glutathione alone cannot reduce insulin.

The data presented in Figure 5 show that all thioredoxins from the parasite can reduce disulphide bridges in insulin, using glutathione as the reducing power. Of the four thioredoxins tested, the highest activity was reported for Trx2 followed by Trx1, Tlp2 and Tlp1. Of note is the fact that Trx2, which exhibits far less activity with TR and NADPH as a reducing system, was efficiently reduced by glutathione. Using glutathione as a reducing system, it showed 4 fold and 5 fold higher activities than Trx1 and Tlp2 respectively. Similarly, Tlp1 which does not serve as a substrate for TR, was effectively reduced by glutathione. Together, these observations lead us to propose that the glutathione and the thioredoxin systems closely interact with each other, and that the glutathione system might reduce those thioredoxins that are poor substrates for thioredoxin reductase (such as Tlp1 and Trx2) or those thioredoxins located at a subcellular compartment lacking thioredoxin reductase (such as Trx2).

**Figure 5.**
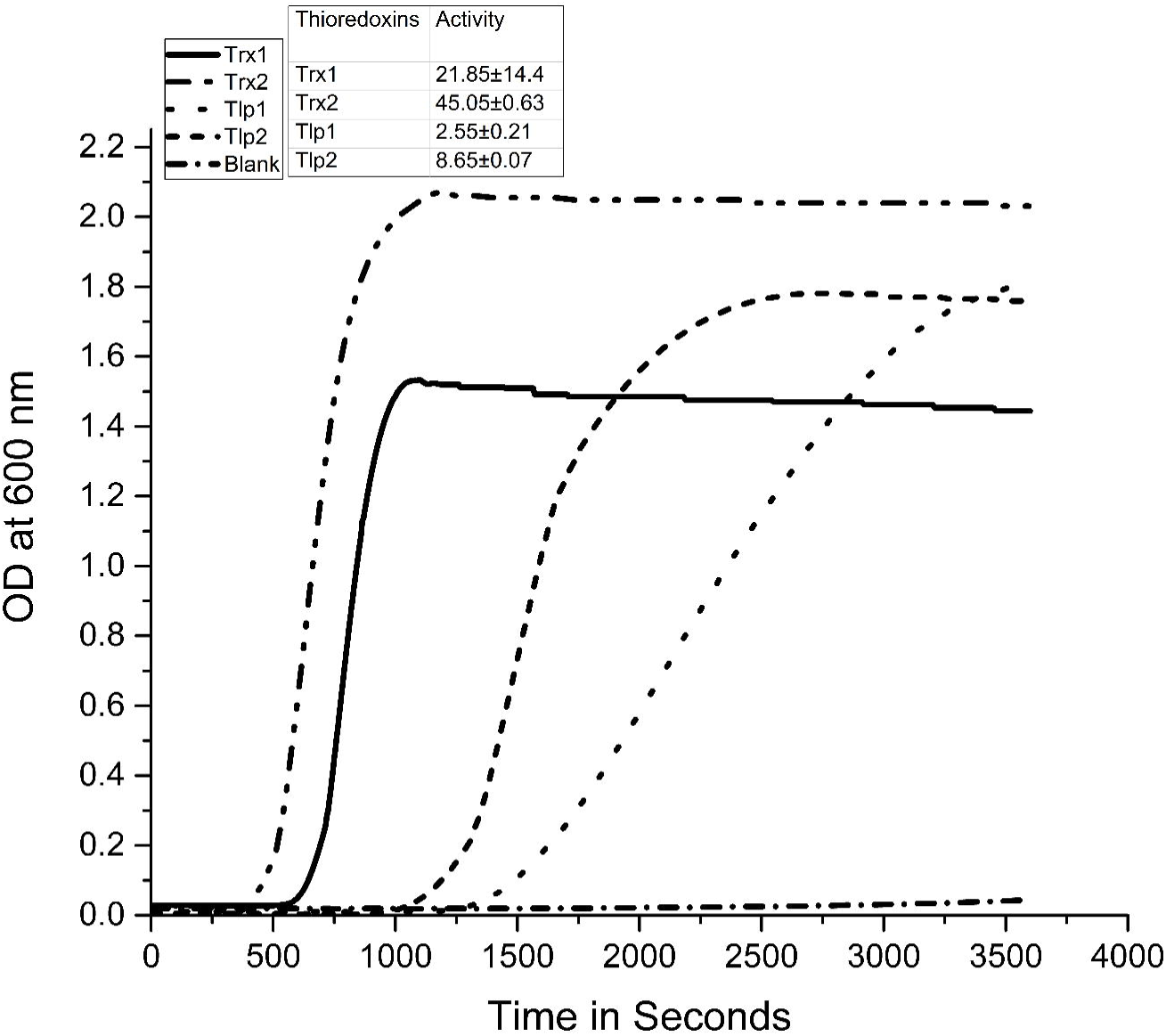
Reduction of *Plasmodium* thioredoxins by glutathione. Turbidity due to thioredoxin mediated insulin precipitation by glutathione was measured at 600nm and plotted as function of time. In this assay protein disulphide reductase activity of thioredoxins should be observed only if thioredoxins are reduced by glutathione. The assay contained 50 mM potassium phosphate buffer (pH 7.4), 1mM EDTA, 170 μM bovine insulin (in 50mM Tris/HCl, 1mM EDTA at pH 7.4), 10mM GSH, 1 U·ml^-1^ S. cerevisiae glutathione reductase (Sigma), 200 μM NADPH and 10μM of respective thioredoxin. The reaction without glutathione served as a negative control. Trx1; thioredoxin 1, Trx2; thioredoxin 2, Trx3, thioredoxin 3, Tlp1; thioredoxin like protein 1, Tlp2; thioredoxin like protein 2. The specific activity of each thioredoxin is shown in inset (expressed as Δ600·min^-2^ × 10^-3^ mg^-1^). The data are the mean ± SD of triplicate reactions.

## Discussion

Antioxidant proteins in the parasite are localized to various subcellular compartments. However, our proposed model network of antioxidant proteins in different compartments of the parasite cell suggested that not all of these compartments are equipped with a complete glutathione or thioredoxin system. These results suggest either that each organelle utilizes only one of the glutathione or thioredoxin systems, or alternatively that the two systems interact with each other. Therefore, we decided to assess the interdependence of the glutathione and thioredoxin systems. We speculated that the absence of a few critical antioxidant proteins from these compartments should have been compensated either by novel proteins or by additional roles played by the existing antioxidant proteins. In this study, we have shown that both glutathione and thioredoxin systems closely interact with each other for the antioxidant defense of the parasite.

The interaction between thioredoxin and glutathione systems has been reported previously in *Plasmodium*. This was, however, with a small subset of proteins. It has been shown that Trx1 from the parasite can support glutathione reduction in a variety of physiological conditions [12]. We performed extensive analyses with multiple thioredoxins that span a majority of the compartments in the parasite. We found that only Trx1 and Tlp2 could support glutathione reduction. This reaction seems to be relevant for those subcellular compartments lacking glutathione reductase, such as the mitochondrion. In support of this hypothesis, our enzymatic analyses using GHOST assays indicate that Tlp2, which is localized to the mitochondrion, can support the reduction of GS-SG, possibly compensating *in vivo* for the lack of glutathione reductase [7]. This phenomenon might be important during the loss of glutathione reductase at a particular stage of the parasite life cycle. For example, *Plasmodium falciparum* merozoites contain glutathione but lack glutathione reductase [13]. Furthermore, a glutathione reductase null mutant of *Plasmodium* is viable in the intra-erythrocytic stage of the life cycle [14]. Therefore, it is necessary that select thioredoxins from the parasite (Trx1 and Tlp2) compensate for this loss of GR function.

An interesting question arises regarding the capacity of the glutathione system to act as a backup in the absence of thioredoxin reductase in a particular subcellular compartment. For instance, the parasitophorous vacuole contains Trx2 as a component of the multimeric PETEX translocon which is responsible for exporting proteins into the host RBCs [15]. Here, Trx2 is speculated to reduce the disulphide bridges thereby unfolding the proteins to facilitate their export through the translocon [15,16]. Therefore, in order to have a functional translocon, Trx2 must be maintained in its reduced state. The parasitophorous vacuole, however, lacks the thioredoxin reductase required for the regeneration of oxidized Trx2, thus suggesting an alternative mechanism of Trx2 reduction [7]. On the other hand, Tlp1 and TR are both localized to the cytosol in the parasite [7]. Although both of these proteins reside in the same compartment, Tlp1 does not get reduced by TR, suggesting the involvement of a different reducing partner. When we evaluated the capacity of *Plasmodium* thioredoxin reductase to serve as an electron donor to Trx2 and Tlp1, we observed that Trx2 exhibited very low activity and Tlp1 did not show any activity. Collectively, these data suggest that there might be alternative mechanisms of thioredoxin reduction in the cell.

It has been recently demonstrated that the glutathione system, particularly glutaredoxin 1 and glutaredoxin 2, can act as backups for the thioredoxin reductase, and have a role in thioredoxin reduction in cells that have lost TrxR activity [17,18]. Recently, it has been shown that, thioredoxin from the parasitic flatworm *Fasciola gigantica* is preferentially reduced by the glutathione system and could be acting as a glutaredoxin [19]. A direct reduction of thioredoxins by glutathione in our biochemical analyses suggests that glutathione might serve as a backup for thioredoxin reductase. In addition to this role, it might be the sole reductant for the thioredoxins that are not exposed to TR (such as Trx2 in parasitophorous vacuole), or for thioredoxins that are not substrates of TR (such as Tlp1 in the cytosol). Based on our results, we speculate that Trx2 and Tlp1 might be reduced by glutathione *in vivo*. Similarly, reduction of thioredoxin by glutathione might also occur at other subcellular compartments where thioredoxin reductase is absent or inactivated by an electrophilic attack.

In our enzymatic analysis, Trx1 and Tlp2 were efficiently reduced by thioredoxin reductase and the localization of these proteins overlaps (Trx1 and TR in cytosol, Tlp2 and TR in mitochondrion). In contrast, Trx2 was found to be a poor substrate for thioredoxin reductase and of note is the fact that, localization of these proteins does not overlap (Trx2 in PV compartment and TR in cytosol). These results show that thioredoxin-thioredoxin reductase pairs can evolve both in function and substrate specificity with respect to their localization. The observed flexibility and adaptations of thioredoxins for their reducing partners due to differential localization makes these proteins particularly suitable for novel redox reactions.

In conclusion, our data suggest that the absence of certain antioxidant proteins in the key subcellular compartments is compensated by crosstalk between the glutathione and thioredoxin systems of the cell. This mutual interaction might strengthen the antioxidant network, thereby ensuring cell survival under constant oxidative stress.

## Acknowledgements

RC thanks Science and Engineering Research Board (SERB) for funding this work (Grant number - SB/YS/LS-354/2013). SP thank Board of Research in Nuclear Sciences (BRNS) for funds (2013/37B/18/BRNS/0489). We thank Aishwarya Narayan and Adhish Walvekar for critically reading the manuscript.

## References

[1] Becker, K., Tilley, L., Vennerstrom, J.L., Roberts, D., Rogerson, S. and Ginsburg, H. (2004). Oxidative stress in malaria parasite-infected erythrocytes: host-parasite interactions. Int J Parasitol 34, 34–163.

[2] Muller, S. (2004). Redox and antioxidant systems of the malaria parasite Plasmodium falciparum. Mol Microbiol 53, 53–1291.

[3] Painter, H.J., Morrisey, J.M., Mather, M.W. and Vaidya, A.B. (2007). Specific role of mitochondrial electron transport in blood-stage Plasmodium falciparum. Nature 446, 88–91.

[4] Ralph, S.A. et al. (2004). Tropical infectious diseases: metabolic maps and functions of the Plasmodium falciparum apicoplast. Nat Rev Microbiol 2, 203–16.

[5] Sato, S., Clough, B., Coates, L. and Wilson, R.J. (2004). Enzymes for heme biosynthesis are found in both the mitochondrion and plastid of the malaria parasite Plasmodium falciparum. Protist 155, 155–117.

[6] van Dooren, G.G., Stimmler, L.M. and McFadden, G.I. (2006). Metabolic maps and functions of the Plasmodium mitochondrion. FEMS Microbiol Rev 30, 596–630.

[7] Kehr, S., Sturm, N., Rahlfs, S., Przyborski, J.M. and Becker, K. (2010). Compartmentation of redox metabolism in malaria parasites. PLoS Pathog 6, e1001242.

[8] Mohring, F., Pretzel, J., Jortzik, E. and Becker, K. (2014). The redox systems of Plasmodium falciparum and Plasmodium vivax: comparison, in silico analyses and inhibitor studies. Curr Med Chem 21, 21–1728.

[9] Aurrecoechea, C. et al. (2009). PlasmoDB: a functional genomic database for malaria parasites. Nucleic Acids Res 37, D539–43.

[10] Holmgren, A. (1979). Thioredoxin catalyzes the reduction of insulin disulfides by dithiothreitol and dihydrolipoamide. J Biol Chem 254, 9627–32.

[11] Martinez-Galisteo, E., Padilla, C.A., Garcia-Alfonso, C., Lopez-Barea, J. and Barcena, J.A. (1993). Purification and properties of bovine thioredoxin system. Biochimie 75, 803–9.

[12] Kanzok, S.M., Schirmer, R.H., Turbachova, I., Iozef, R. and Becker, K. (2000). The thioredoxin system of the malaria parasite Plasmodium falciparum. Glutathione reduction revisited. J Biol Chem 275, 40180–6.

[13] Krauth-Siegel, R.L., Muller, J.G., Lottspeich, F. and Schirmer, R.H. (1996). Glutathione reductase and glutamate dehydrogenase of Plasmodium falciparum, the causative agent of tropical malaria. Eur J Biochem 235, 345–50.

[14] Pastrana-Mena, R. et al. (2010). Glutathione reductase-null malaria parasites have normal blood stage growth but arrest during development in the mosquito. J Biol Chem 285, 27045–56.

[15] de Koning-Ward, T.F. et al. (2009). A newly discovered protein export machine in malaria parasites. Nature 459, 945–9.

[16] Matthews, K. et al. (2013). The Plasmodium translocon of exported proteins (PTEX) component thioredoxin-2 is important for maintaining normal blood-stage growth. Mol Microbiol 89, 1167–86.

[17] Du, Y., Zhang, H., Lu, J. and Holmgren, A. (2012). Glutathione and glutaredoxin act as a backup of human thioredoxin reductase 1 to reduce thioredoxin 1 preventing cell death by aurothioglucose. J Biol Chem 287, 38210–9.

[18] Zhang, H., Du, Y., Zhang, X., Lu, J. and Holmgren, A. (2014). Glutaredoxin 2 reduces both thioredoxin 2 and thioredoxin 1 and protects cells from apoptosis induced by auranofin and 4-hydroxynonenal. Antioxid Redox Signal 21, 669–81.

[19] Gupta, A., Pandey, T., Kumar, B. and Tripathi, T. (2015). Preferential regeneration of thioredoxin from parasitic flatworm Fasciola gigantica using glutathione system. Int J Biol Macromol 81, 983–90.

